# The impact of Temporal Artery Biopsy for the diagnosis of Giant Cell Arteritis in clinical practice in a tertiary University Hospital

**DOI:** 10.1101/512137

**Authors:** E Kaltsonoudis, E Pelechas, A Papoudou-Bai, E.T. Markatseli, M Elisaf, PV Voulgari, AA Drosos

**Affiliations:** Rheumatology Clinic, Department of Internal Medicine, Medical School, University of Ioannina, Greece; Department of Pathology, Medical School, University of Ioannina, Greece; Department of Internal Medicine, Medical School, University of Ioannina, Greece

**Keywords:** Giant cell arteritis, temporal artery biopsy, polymyalgia rheumatica, visual disturbances, fever of unknown origin, new headache

## Abstract

**Background:** Temporal artery biopsy (TAB) is useful in assisting with giant cell arteritis (GCA) diagnosis but lacks sensitivity. The aim of our study was to assess the diagnostic impact of TAB histology in patients with suspected GCA on hospital admission.

**Methods:** A prospectively maintained database was queried for all TABs performed between 1-1-2000 until 31-12-2017 at the University Hospital of Ioannina. Thus, inclusion criteria were made on the grounds of every patient that underwent a TAB during the above-mentioned period, regardless of demographic, clinical and laboratory data.

**Results:** Two hundred forty-five TABs were included (149 females and 96 males), with a mean age of 64.5 (±3.5) years. The mean symptoms duration until admission to the hospital was 8.6 (±1.3) weeks and all had elevated acute phase reactants on admission. The reasons of admission were fever of unknown origin (FUO) in 114 (46.5%) patients, symptoms of polymyalgia rheumatica (PMR) in 84 (34.3%), new headache in 33 (13.5%), anemia of chronic disease (ACD) in 8 (3.32%) and eye disturbances in 6 (2.5%) patients. Positive results were found in 49 (20%) TABs. More specifically, in 14% of patients with FUO, 21% in those with PMR, while in patients with a new headache the percentage was 27%. Finally, 5 out of 6 (83.3%) of patients with ocular symptoms and only one (12.5%) of those suffering from ACD. Visual manifestations and FUO are correlated with a positive TAB.

**Conclusion:** It seems that TAB is useful in assisting with GCA diagnosis, but lacks sensitivity.

## INTRODUCTION

Giant cell arteritis (GCA), also referred to as cranial arteritis, temporal arteritis or Horton’s disease is a type of systemic inflammatory vasculitis of unknown etiology. It is the commonest form of vasculitis in the elderly [1], which, if left untreated, may cause blindness [2] and stroke [3]. Even if it is classified as a large-vessel vasculitis (LVV), after the 2012 revised International Chapel Hill Consensus Conference, medium and small arteries are also involved [4]. Typically, it affects the superficial temporal arteries (hence the term temporal arteritis), the ophthalmic, occipital and vertebral arteries but also the aorta, carotid and subclavian arteries. The inflammation leads to vessel functional impairment (stenoses), and as a consequence, to diminished tissue blood supply. Subsequently, irreversible damage may develop affecting mainly the eyes and the central nervous system. It is more prevalent in people over the age of 50 [5].

The diagnosis of GCA is based on the 1990 ACR criteria which requires 3 out of 5 of these to be present [6]. However, accurate early GCA diagnosis can be established through temporal artery biopsy (TAB), which is the gold standard method [7]. In addition, imaging techniques like computed tomography (CT), CT angiography, magnetic resonance (MR), MR angiography, color–Doppler sonography (CDS) and high-resolution CDS, as well as positron emission tomography/computed tomography (PET/CT) can be used [8–10]. On the other hand, while the non-invasive imaging techniques offer information regarding the diagnosis and treatment, the histological confirmation by TAB allows an objective documentation of the diagnosis. To this end, an appropriate therapeutic strategy may be crucial regarding the evolution of the disease. Furthermore, whenever required, a more rational aggressive and personalized therapy with steroids or with the newer and promising agents can be applied [11].

The aim of our study was to assess the impact of TAB histology on the clinical diagnosis in patients who were admitted in the hospital, and the suspected diagnosis of GCA should be excluded.

## MATERIAL AND METHODS

A prospectively maintained database was queried for all TABs performed between 1-1-2000 until 31-12-2017 at the University Hospital of Ioannina. All patients’ records were analyzed for demographic, clinical and laboratory data undergoing a TAB, during the period of January 2000 to December 2017. Thus, inclusion criteria were: any patient regardless of demographic, clinical and laboratory data undergoing a TAB during the above-mentioned period. All TABs were carried out prior to steroid therapy. Statistical analysis was performed using SPSS Statistics, version 20.0. An informed consent form has been obtained by all patients and the study has been approved by the hospital’s ethics committee.

### Definitions

***Fever of unknown origin (FUO):*** is a disease condition of temperature exceeding 38.3°C on at least three occasions over a period of at least three weeks, with no diagnosis made, despite one week of inpatient investigation [12].

***Polymyalgia Rheumatica (PMR):*** is defined when a patient complains for pain or/and stiffness affecting the neck, shoulders and hips in individuals over 50 years of age.

***New headache:*** is a new onset headache which is not caused by another known condition or disorder.

***Anemia of chronic disease (ACD):*** normocytic, normochromic anemia with low serum iron and normal or high serum ferritin levels.

***Visual disturbances:*** defined as acute eye pain, diplopia, amaurosis fugax or visual loss of varying severity.

***Positive TAB:*** when the histological findings of the temporal artery showed inflammatory infiltrates of the arterial wall, with or without the presence of giant cells and/or rupture of the internal elastic lamina.

## RESULTS

Two hundred forty-five TABs were included. There were 149 females and 96 males with a mean age of 64.5 (±3.5) years, who underwent TABs. The mean symptoms duration until admission to the hospital was 8.6 (±1.3) weeks and all had elevated erythrocyte sedimentation rate (ESR) or/and C-reactive protein (CRP). Figure shows that from 245 TABs, positive results were found in 49 (20%). Twelve biopsies samples were insufficient to confirm or refute GCA diagnosis. This figure depicts also the departments ordered the TABs, as well as the clinical features at admission and the positive TAB histology achieved by each clinic. Positive TABs ranged from zero, ordered by the orthopedic clinic to 83.3% ordered by the eye clinic. The rheumatology clinic achieved positive TABs in 23% of the cases and the department of internal medicine 19%. Finally, 8.3% positive TABs achieved by the neurology clinic. The demographic, clinical, laboratory and histological data are presented in the Table 1. Half of the patients were admitted due to FUO. In this group positive TABs were demonstrated in 14%. One third of patients was admitted with symptoms of PMR. In these patients positive TABs were found in 21%, while in patients with new headache positive TABs were demonstrated in 27%. Finally, 5 out of 6 (83.3%) of patients admitted with ocular symptoms had positive TABs and only 1 (12.5%) from patients suffering from ACD had positive TAB. Table 2 shows that visual manifestations and FUO are correlated with positive TAB.

**Figure:**
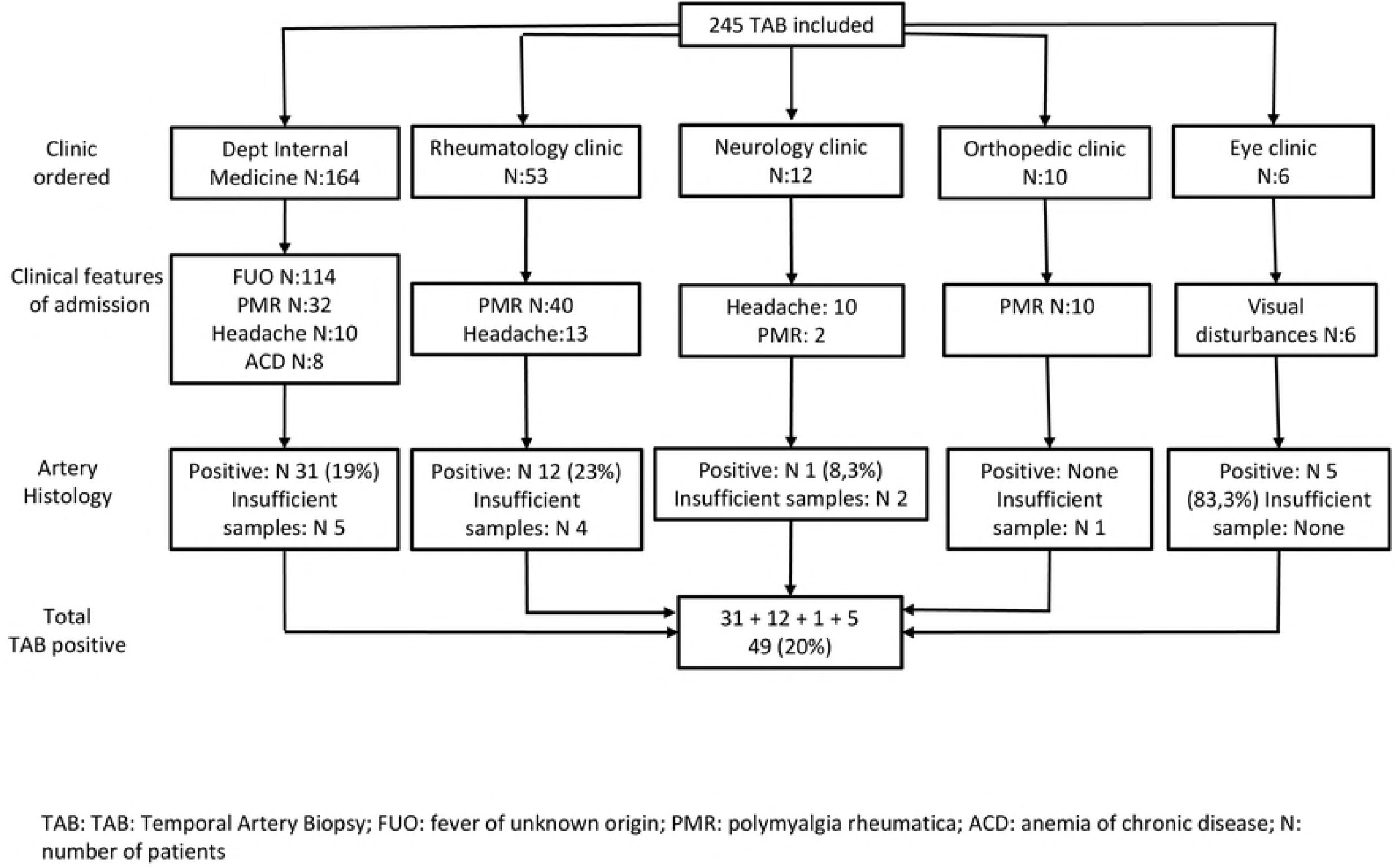
Diagrammatic representation of TABs ordered by different clinics during admission

**Table 1.** Demographic, clinical and laboratory data of patients with suspected GCA who underwent TABs.

| Parameters | Values | Positive TAB n (%) |
| --- | --- | --- |
| Total number of patients | 245 | 49 (20) |
| Female/male | 149/96 | 30/19 (20/18) |
| Mean age (± years) | 64.5 (3.5) | |
| Mean symptoms duration until admission (± weeks) | 8.6 (1.3) | |
| Clinical symptoms and signs |  |  |
| • Fever of unknown origin n (%) | 114 (46.5) | 16 (14) |
| • Polymyalgia rheumatica n (%) | 84 (34.3) | 18 (21) |
| • New headache n (%) | 33 (13.5) | 9 (27) |
| • Anemia of chronic disease | 8 (3.2) | 1 (12.5) |
| • Visual disturbance n (%) | 6 (2.5) | 5 (83.3) |
| Elevated acute phase reaction |  |  |
| • ESR n (%) | 245 (100) |  |
| • CRP n (%) | 215 (84) |  |
| • ESR + CRP n (%) | 215 (84) |  |

GCA: Giant Cell Arteritis; TAB: Temporal Artery Biopsy; ESR: Erythrocyte Sedimentation Rate; CRP:
C-reactive protein;

**Table 2.**
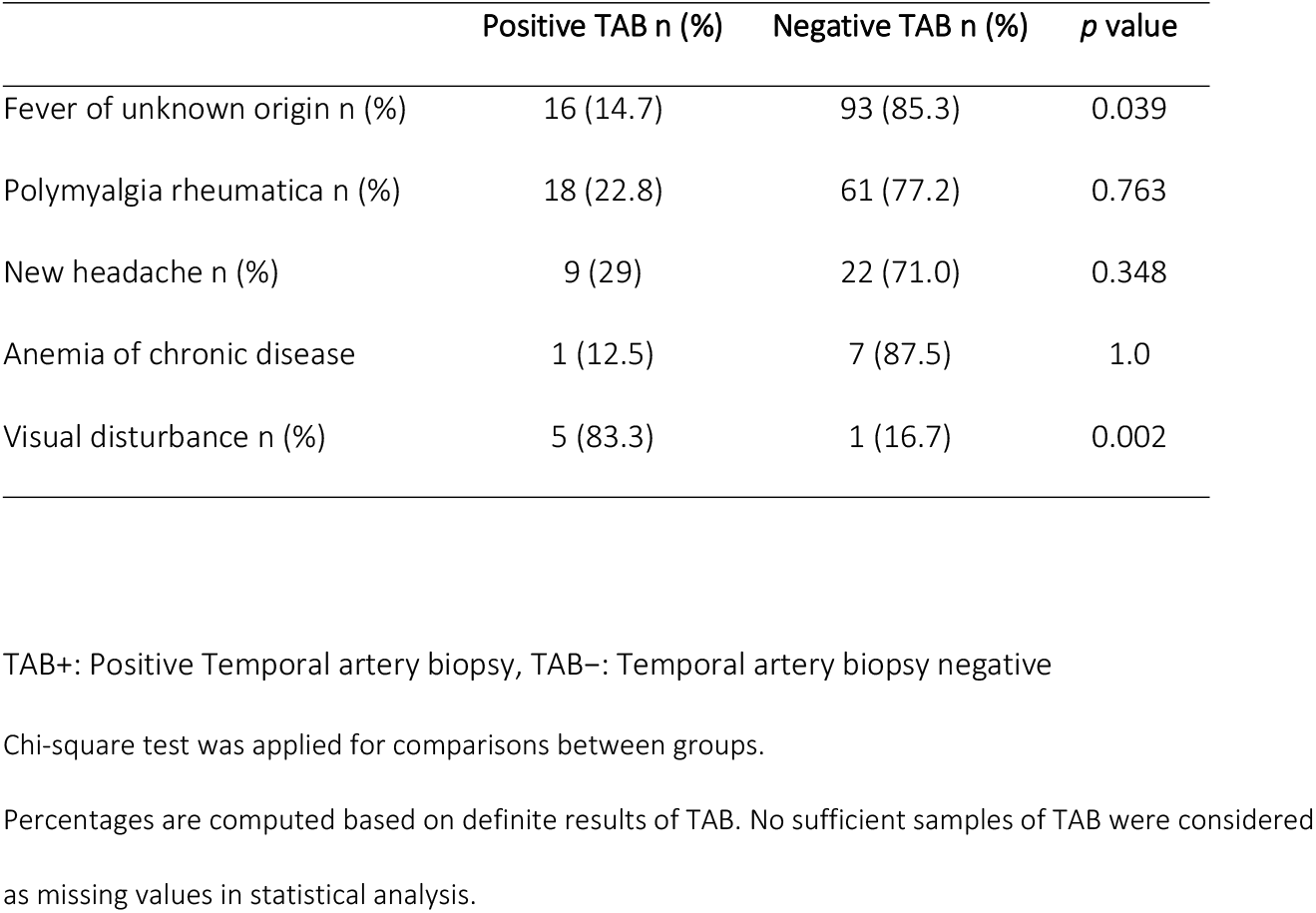
Clinical variables in TAB+ and TAB-groups.

|  | Positive TAB n (%) | Negative TAB n (%) | *p* value |
| --- | --- | --- | --- |
| Fever of unknown origin n (%) | 16 (14.7) | 93 (85.3) | 0.039 |
| Polymyalgia rheumatica n (%) | 18 (22.8) | 61 (77.2) | 0.763 |
| New headache n (%) | 9 (29) | 22 (71.0) | 0.348 |
| Anemia of chronic disease | 1 (12.5) | 7 (87.5) | 1.0 |
| Visual disturbance n (%) | 5 (83.3) | 1 (16.7) | 0.002 |

TAB+: Positive Temporal artery biopsy, TAB–: Temporal artery biopsy negative
Chi-square test was applied for comparisons between groups.
Percentages are computed based on definite results of TAB. No sufficient samples of TAB were considered as missing values in statistical analysis.

## DISCUSSION

Currently there is no 100% accurate test for GCA. Elderly patients usually have a new-onset headache and scalp tenderness, typically with an abnormal laboratory test, usually high ESR. Applying the ACR criteria for GCA, these patients are classified as having temporal arteritis. However, it can be difficult to distinguish non-serious forms of headache from GCA. Infections produce similar clinical signs and abnormal laboratory tests. If there is a strong suspicion of GCA, treatment with steroids should be initiated immediately. To confirm the diagnosis, the patient should undergo a biopsy of the temporal artery. On the other hand, biopsy of the temporal artery for the diagnosis of GCA has a relatively low yield [13]. The difficulty in diagnosing GCA, is the lack of a high-quality gold standard test. Although biopsy is reported to be the current gold standard test for its diagnosis, the majority of patients in whom a diagnosis of GCA in suspected do not actually have a positive result [13]. This may reflect the fact that there is a higher index of suspicion for diagnosis and therefore more patients with headache are being evaluated for GCA. Equally, it may reflect the relatively poor association between the true multi-vessel disease of GCA and the TAB findings to support a diagnosis of GCA.

In the present study, overall, the number of positive biopsies remains low (20%). A possible explanation could be that the clinical picture of GCA has a wide spectrum of manifestations ranging from cranial manifestations (headache, scalp tenderness, jaw claudication, visual disturbances), to PMR symptoms or to constitutional symptoms with weakness low grade fever, arthralgias, or symptoms affecting the large vessels (LV) [14]. In addition, TAB biopsies have been ordered by various clinics and specialties which included an unselected population influencing TAB positivity. The involvement of LV is more than 40% of GCA patients and in this case half of the patients have negative TAB. On the other hand, in patients with PMR, TAB biopsy is positive in about 20%. Indeed, as we described in the Table 1, the majority of TABs were ordered by the Department of Internal Medicine in which patients were admitted with symptoms of FUO, PMR or headache. The majority of them had negative TABs. In this case, TABs were ordered to rule out GCA, since it is included as a cause of FUO. In contrast, 5 out of 6 (83,3%) of TABs ordered by the eye clinic were positive. It seems that when a TAB is performed in patients with symptomatology of cranial involvement versus those with other clinical manifestations, there is a higher yield of a positive result for the former. Thus, jaw claudication and blindness are reported as predictive variables for a positive TAB while headaches and PMR are associated with lower rates of positivity. However, in a multivariate analysis these variables do not reach statistical significance [15].

Imaging modalities have been studied extensively as potential diagnostic markers for GCA. Ultrasound (US) with CDS is the most practical and widely used modality for investigating suspected GCA [16–18]. Three meta-analyses have supported the role of US in the diagnosis of GCA [19–21]. The presence of bilateral US abnormalities (both temporal arteries involved) provides high specificity (100%) for the diagnosis of GCA, but it is sensitivity was 43%. Two of the meta-analyses, reported concerns with the quality of the included studies [21,22] and the third did not access the methodology (19). Currently, the use of US as a diagnostic tool for GCA is relatively limited, perhaps as a result of practical reasons relating to training how to use the US or equipment availability to facilitate rapid access and evaluation of patients with suspected GCA. On the other hand, US examination of temporal arteries is not invasive and there is no ionizing radiation involved. Furthermore, it can provide information about the entire length of the temporal arteries. In addition, examination of the axillary arteries improves the sensitivity of US for the diagnosis of GCA showing a high sensitivity (54 vs 39%) but lower specificity (81 vs 100%) for US compared to TAB in diagnosing GCA [22].

Published data clearly suggest a positive value of PET/CT in securing the diagnosis of LVV in treating naïve patients. A meta-analysis showed that PET had a sensitivity of 90% and specificity of 98% for the diagnosis of GCA [23]. PET is valuable for diagnostic purpose in those patients who present with atypical manifestations, for examples in patients with GCA without cranial symptoms and showed to increase the diagnostic accuracy over the above standard workup examination [23, 24]. However, PET is less available and is considered more expensive than CT or MR. A limitation of this report is that our study is a retrospective one, however it includes a large number of patients who underwent TABs from different specialties and clinics and this represent a real-life work-up in a tertiary University Hospital. On the other hand, our results are in line with reports from Spain and France suggesting that elderly patients mainly with visual manifestations and jaw claudication have positive TABs [25,26]. Our observations confirm the presence of different disease patterns of clinical presentation in GCA (14, 27) and emphasize the importance of eye manifestations and FUO as factors that may predict TAB positivity.

In summary, the management of GCA requires a balance between ensuring that patients with GCA are diagnosed and treated appropriately and avoiding the burden of unnecessary steroids treatment. Not all elderly patients with a new-onset headache and a high ESR had GCA, despite the fact that they satisfied the ACR criteria. Thus, the documentation of GCA diagnosis is an imperative. TAB is a useful procedure that helps in this direction, but unfortunately, it lacks sensitivity. On the other hand, both US and PET scans, have a role as an alternative to or in addition to biopsy for GCA diagnosis. However, the routine use of US and PET scan for GCA is restricted to only a few centers. Thus, TAB remains the standard test for the majority of patients suspected as having GCA.

## AUTHOR CONTRIBUTIONS

All authors researched data for the article, contributed substantially to discussions of its content and wrote the manuscript.

A.A.D. and P.V.V. contributed to review and editing of the article before submission.

## Abbreviations

GCA: Giant cell arteritis
LVV: large-vessel vasculitis
CT: computed tomography
MR: magnetic resonance
CDS: color–Doppler sonography
PET/CT: positron emission tomography/computed tomography
FUO: Fever of unknown origin
PMR: Polymyalgia Rheumatica
ACD: Anemia of chronic disease
ESR: erythrocyte sedimentation rate
CRP: C-reactive protein
LV: large vessels
US: Ultrasound

